# Role of necroptosis in chronic hepatic inflammation and liver disease in Cu/Zn superoxide dismutase deficient mice

**DOI:** 10.1101/2020.08.17.254888

**Authors:** Sabira Mohammed, Evan H Nicklas, Nidheesh Thadathil, Ramasamy Selvarani, Gordon H Royce, Arlan Richardson, Sathyaseelan S. Deepa

## Abstract

Mice deficient in the antioxidant enzyme Cu/Zn-superoxide dismutase (*Sod1*^−/−^ or Sod1KO mice) develop spontaneous hepatocellular carcinoma (HCC) with age. Similar to humans, HCC development in Sod1KO mice progresses from fatty liver disease to non-alcoholic steatohepatitis (NASH) with fibrosis, which eventually progresses to HCC. Because liver inflammation is the main mechanism that drives the disease progression in chronic liver disease (CLD) and because necroptosis is a major source of inflammation, we tested the hypothesis that increased necroptosis in the liver plays a role in increased inflammation and progression to fibrosis and HCC in Sod1KO mice. Phosphorylation of MLKL (P-MLKL), a well-accepted marker of necroptosis, and expression of MLKL protein were significantly increased in the livers of Sod1KO mice compared to WT mice indicating increased necroptosis. Similarly, phosphorylation of RIPK3 and RIPK3 protein levels were also significantly increased. Markers of pro-inflammatory M1 macrophages, NLRP3 inflammasome, and transcript levels of pro-inflammatory cytokines and chemokines, e.g., TNFα, IL-6, IL-1β, and Ccl2 that are associated with human NASH and HCC, were significantly increased. Markers of fibrosis and oncogenic transcription factor STAT3 were also upregulated in the livers of Sod1KO mice. Short term treatment of Sod1KO mice with necrostatin-1s (Nec-1s), a necroptosis inhibitor, significantly reduced necroptosis, pro-inflammatory cytokines, fibrosis markers and STAT3 activation. Our data show for the first time that necroptosis-mediated inflammation contributes to fibrosis and HCC progression in Sod1KO mice, a mouse model of accelerated aging and progressive HCC development. These findings suggest that necroptosis might be a target for treating NASH and HCC.

## Introduction

Chronic liver disease (CLD) is a major health problem affecting over 800 million people worldwide with an estimated mortality rate of 2 million deaths per year (1,2). CLD involves a spectrum of diseases ranging from viral infection (hepatitis B and C) to alcohol induced CLD and metabolic CLD such as non-alcoholic fatty liver disease (NAFLD) and non-alcoholic steatohepatitis (NASH). Due to the obesity pandemic in recent years, the increased incidence of NAFLD affecting nearly 25% of the world population has become one of the leading risk factors for CLD-associated mortality (3). NAFLD arising from obesity can eventually progress to non-alcoholic steatohepatitis (NASH), a condition characterized by increased steatosis, liver inflammation and hepatocellular damage. Nearly one-third of patients with NASH develop advanced fibrosis or cirrhosis that can progress to hepatocellular carcinoma (HCC) (4, 5), which is the fastest-growing cause of cancer-related deaths in the United States and world (6). Liver inflammation has been identified as the one of the major mechanisms driving the progression of CLD and HCC(7).

Several factors have been identified that trigger liver inflammation in CLD, including extrinsic factors derived from gut or adipose tissue and intrinsic factors in liver, e.g. innate immune response, lipotoxicity and cell death pathways (8). Of these, hepatocyte cell death is one of the key factors that can cause liver inflammation in CLD (9). Several forms of hepatocyte cell death have been reported in CLD, e.g., necrosis, apoptosis, pyroptosis and necroptosis (programmed necrosis). Whereas apoptosis is considered as a non-inflammatory mode of cell death, several studies have demonstrated that necroptosis plays a major role in inflammation (10–12). Necroptotic stimuli (e.g., TNF-α, oxidative stress, and mTOR/Akt signaling) sequentially activate the three key kinases in the necroptosis pathway through phosphorylation: receptor-interacting serine/threonine-protein kinase 1 (RIPK1), RIPK3 and mixed lineage kinase domain like pseudokinase (MLKL). Phosphorylation of MLKL leads to its oligomerization and binding to the plasma membrane leading to the disruption of the membrane and the release of cellular components including damage associated molecular patterns (DAMPs) (e.g., mitochondrial DNA, S100A9, HMGB-1, ATP, IL-33 etc.). DAMPs initiate and exacerbate the inflammatory process by binding to cell surface receptors of innate immune cells(13, 14). Studies have shown that necroptosis is increased in NAFLD/NASH in mice, and inflammation associated with NAFLD/NASH with high fat feeding in mice is reduced by inhibiting necroptosis (15–21). In addition, studies with human subjects show that increased necroptosis is associated with NASH (16, 17).

To study the role of necroptosis in CLD including HCC, we used a novel mouse model that develops spontaneous NAFLD/NASH and progresses to HCC with age: mice deficient in the antioxidant enzyme, Cu/Zn superoxide dismutase (*Sod1*^−/−^ or Sod1KO mice). Sod1KO mice show increased accumulation of triglyceride in the liver, a characteristic feature of NAFLD, as early as 3 weeks of age (22). The Sod1KO mice also show increased accumulation of collagen in liver indicating increased fibrosis (23) and develop spontaneous HCC around 18 to 20 months of age (24). The Sod1KO mice have several features that make them particularly relevant to the progression of CLD and HCC in humans. First, although they do not become obese, the Sod1 KO mice develop fatty liver from an imbalance in triglyceride deposition and removal arising from impaired very low density lipoprotein (VLDL) secretion because apolipoprotein B (ApoB) levels are reduced in the livers of Sod1KO mice (22), and APOB mutations are associated with HCC in humans (25). Second, the increased oxidative stress associated with knocking out *Sod1* is a key factor in the development of HCC in the Sod1KO mice (24, 26), and oxidative stress is one of the factors that has been proposed to play a role in HCC in humans (27). Third, HCC, like most other cancers, is age dependent with the peak incidence of HCC occurring after 65 years of age. All of the current HCC mouse models exhibit HCC in relatively young adult mice. The Sod1KO mice, which show an accelerated aging phenotype (28), develop HCC in the later third of their life, similar to what is seen in humans. Because a variety of factors change with age that can impact cancer, the Sod1KO mice allows one to study HCC in the environment of an aged animal that is relevant to HCC in humans.

Previously we reported that necroptosis and expression of proinflammatory cytokines are increased in the adipose tissue of Sod1KO mice (28). Therefore, we hypothesized that increased necroptosis in the liver plays a role in increased inflammation and progression to fibrosis and HCC reported in Sod1KO mice. To test this hypothesis, we treated Sod1KO mice with a necroptosis inhibitor, necrostatin-1s (Nec-1s), which targets RIPK1 (29) and has been shown to block necroptosis and reduce expression of proinflammatory cytokines in mouse models of multiple sclerosis and amyotrophic lateral sclerosis (30,31). Our findings show that markers of necroptosis, inflammation and fibrosis are increased, and oncogenic transcription factor STAT3 is activated in the livers of Sod1KO mice. Importantly, we show that Nec-1s treatment reduced markers of necroptosis, inflammation (levels of pro-inflammatory cytokines and chemokines, M1 macrophage markers), fibrosis and STAT3 activation in the livers of Sod1KO mice.

## Results

In these experiments, we tested whether necroptosis and proinflammatory cytokines are increased in the livers of Sod1KO mice and whether blocking necroptosis has any effect on the expression of proinflammatory cytokines. To block necroptosis, we treated the mice with Nec-1s for 25 days, which inhibits RIPK1 and has been shown to effectively block necroptosis and reduce expression of proinflammatory cytokines in multiple sclerosis (30) and amyotrophic lateral sclerosis (31). Three groups of 10 to 13-month-old male mice were used for the study: (1) WT mice (vehicle treated), (2) Sod1KO mice (vehicle treated), and (3) Sod1KO mice (Nec-1s treated). Because Sod1KO mice are reported to have reduced body weight and increased liver weight compared to control mice (26), we assessed the effect of Nec-1s treatment of these parameters. We found that Nec-1s treatment had no effect on body weight or liver weight of Sod1KO mice (Supplementary Figures 1A and 1B). Thus Nec-1s treatment had no obvious negative effect on Sod1KO mice.

### Effect of Sod1 deficiency on markers of necroptosis

Phosphorylation of MLKL (P-MLKL) is a well-accepted marker of necroptosis because the membrane binding of P-MLKL is the key step in disrupting the cell membrane (32). Therefore, we first tested whether necroptosis is activated in the livers of Sod1KO mice by measuring the levels of P-MLKL and MLKL. As shown in Figure 1A, the levels of both P-MLKL and MLKL were significantly increased (1.7-and 5.8-fold, respectively) in the livers of Sod1KO mice compared to WT mice indicating increased necroptosis. Nec-1s treatment resulted in a significant reduction in the expression of P-MLKL and MLKL in the livers of Sod1KO mice, such that P-MLKL levels were not significantly different from the WT control mice (Figure 1A). Next, we assessed changes in the expression and phosphorylation of other proteins involved in the necroptosis pathway, e.g., RIPK3 and RIPK1. Necroptosis is initiated by the phosphorylation of RIPK1 that catalyzes the phosphorylation of RIPK3, which in turn phosphorylates MLKL leading to membrane rapture. Phosphorylation of RIPK3 (P-RIPK3) and RIPK3 protein levels were significantly increased (2.5-fold and 2-fold, respectively) in Sod1KO mice liver, and Nec-1s treatment significantly reduced the levels of P-RIPK3 and RIPK3 (Figure 1B). There was no significant change in the phosphorylation of RIPK1 in Sod1KO mice liver compared to WT mice and Nec-1s treatment had no effect on the levels of P-RIPK1. The levels of RIPK1 protein were significantly lower in Sod1KO mice liver compared to WT mice, and Nec-1s did not alter RIPK1 protein expression (Figure 1B). Figure 1C shows the effect of Nec-1s on the transcript levels of RIPK1, RIPK3 and MLKL. Transcript levels of RIPK3 (1.8-fold) and MLKL (4-fold), were significantly increased in Sod1KO mice liver compared to WT mice, and Nec-1s resulted in a significant reduction in the expression of RIPK3 and MLKL in Sod1KO mice liver. The levels of RIPK1 transcripts were similar in WT and Sod1KO mice liver, and Nec-1s treatment had no effect on the expression of RIPK1 in Sod1KO mice liver (Figure 1C). Thus, the RIPK3 and MLKL components of the necroptosis pathway are increased in the livers of Sod1KO mice largely through an increase in transcription, and Nec-1s effectively blocked the increase in the expression of these components of necroptosis.

**Figure 1.**
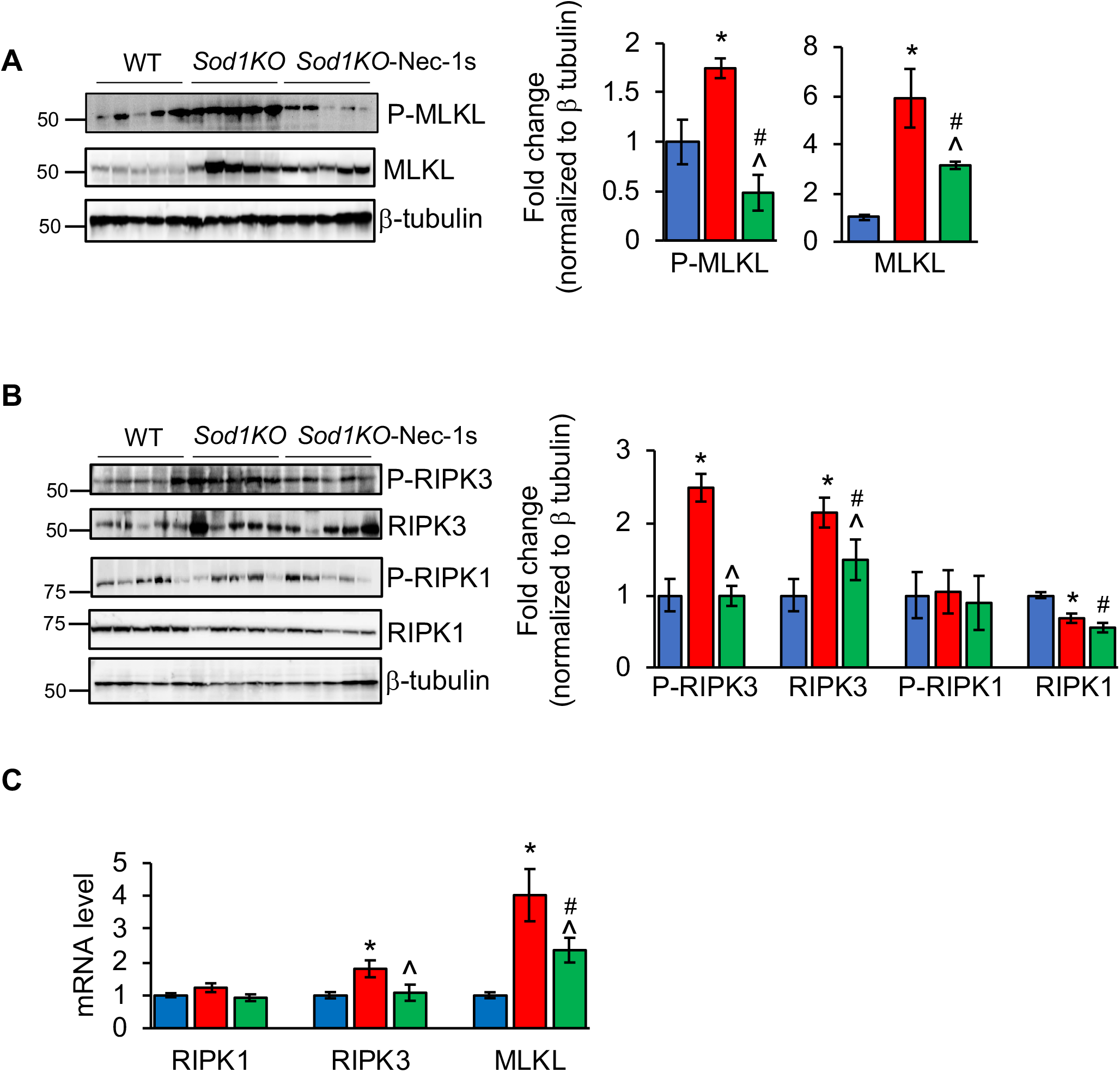
Effect of *Sod1* deficiency on necroptosis. **(A)** *Left panel:* Immunoblots of liver extracts prepared from WT (blue bars), Sod1KO untreated (red bars) and Sod1KO mice treated with Nec-1s (green bars) for P-MLKL, MLKL and β-tubulin. *Right panel*: Graphical representation of quantified blots normalized to β-tubulin. **(B)** *Left panel:* Immunoblots of liver extracts for P-RIPK3, RIPK3, P-RIPK1, RIPK1 and β-tubulin. *Right panel*: Graphical representation of quantified blots normalized to β-tubulin. **(C)** Transcript levels for MLKL, RIPK3, and RIPK1. Data were obtained from 5 to 7 mice per group and are expressed as the mean + SEM. (ANOVA, *WT-Veh vs Sod1KO-Veh; ^#^WT-Veh vs Sod1KO-Veh; ^^^ Sod1KO-Veh vs Sod1KO-Nec-1s; ^*/#/^^ P ≤ 0.05).

Because RIPK3 protein is involved in the activation of both apoptosis and necroptosis, we tested whether apoptosis was activated in Sod1KO mice liver and whether blocking necroptosis had any effect on apoptosis. Apoptosis was assessed by measuring the level of cleaved caspase-3, the active form of caspase-3. Levels of cleaved caspase-3 normalized to caspase-3 was significantly increased in Sod1KO mice liver compared to WT mice and Nec-1s had no effect on the levels of cleaved caspase-3 in Sod1KO mice liver (Supplementary Figure 1C). Thus, both apoptosis and necroptosis are activated in Sod1KO mice liver. However, Nec-1s treatment attenuated only necroptosis; it had no effect on apoptosis in the livers of Sod1KO mice.

### Effect of Sod1 deficiency and necroptosis on inflammation

Because DAMPs released by necroptosis are key in the activation of the inflammatory response, and macrophages are involved in the production of inflammatory cytokines, the expression of F4/80, a marker of mice liver macrophages (Kupffer cells) was assessed. A significant increase in the transcript levels of macrophage marker F4/80 (7.4-fold) was observed, and Nec-1s treatment significantly reduced their expression (Figure 2A). Macrophages are categorized into M1 or M2 phenotypes. In general, M1 macrophages play a more proinflammatory role in liver injury and M2 macrophages exert an anti-inflammatory effect. In Sod1KO mice liver, markers of M1 macrophages CD68 (5.4-fold), CD86 (4.6-fold), and TLR4 (5.5-fold) were significantly elevated compared to WT mice, and Nec-1s treatment significantly reduced their expression (Figure 2B). Markers of M2 macrophages were either down regulated (Arg1) or unaltered (CD206) in Sod1KO mice liver. Nec-1s treatment had no effect on the expression of Arg1 or CD206 (Supplementary Figure 2A).

**Figure 2.**
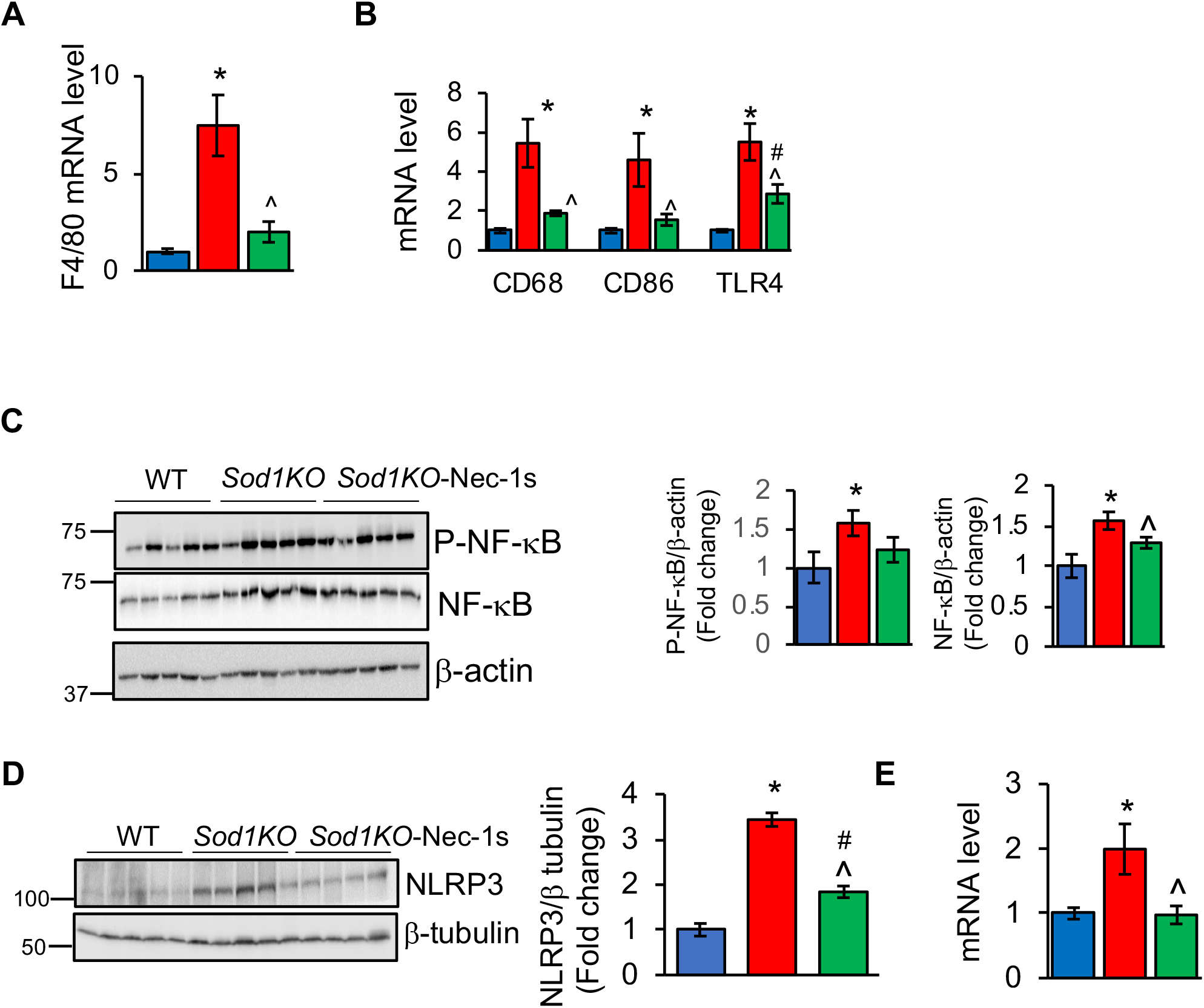
Effect of *Sod1* deficiency and necroptosis on inflammation. Transcript levels of F4/80 **(A)**, and proinflammatory M1 macrophage markers CD68,CD86, and TLR4 **(B)** in the livers of WT (blue bars), Sod1KO untreated (red bars) and Sod1KO mice treated with Nec-1s (green bars). **(C)** *Left panel:* Immunoblots of liver extracts for phospho-NF-κB (p65), NF-κB and β-actin. *Right panel*: Graphical representation of quantified blots normalized to β-actin. **(D)** *Left panel:* Immunoblots of liver extracts for NLRP3 and β-tubulin. *Right panel*: Graphical representation of quantified blots normalized to β-tubulin. **(E)** Transcript levels of NLRP3. Data were obtained from 5 to 7 mice per group and are expressed as the mean + SEM. (ANOVA, *WT-Veh vs Sod1KO-Veh; ^#^WT-Veh vs Sod1KO-Veh; ^^^ Sod1KO-Veh vs Sod1KO-Nec-1s; ^*/#/^^ P ≤ 0.05).

Activation of the NF-κB pathway in macrophages by DAMPs is one of mechanisms proposed for increased production of pro-inflammatory cytokines (33). Therefore, we determined whether NF-κB pathway is activated in the livers of Sod1KO mice by assessing the expression of active form of NF-κB, phospho-NF-κB p65. The levels of phospho-NF-κB p65 were significantly increased in the livers Sod1KO mice (1.5-fold) compared to WT mice, and Nec-1s treatment showed a tendency to reduce the levels of phospho-NF-κB p65; however, this decrease was not statistically significant. The levels of NF-κB protein were also significantly increased in Sod1KO mice liver (1.5-fold) and was significantly reduced by Nec-1s treatment (Figure 2C).

DAMPs activate the NLRP3 inflammasome complex in macrophages, which leads to the secretion of proinflammatory cytokines such as IL-1β, IL-18 (34). Therefore, we measured the levels of NLRP3 (NOD-, LRR- and pyrin domain-containing protein 3), which is a component of the inflammasome. NLRP3 protein and transcript levels were significantly increased (3.4-fold) in the livers of Sod1KO mice compared to WT mice, and Nec-1s treatment significantly reduced NLRP3 protein and transcript levels in the livers of Sod1KO mice to levels similar to the WT control mice (Figures 2D and 2E).

We next measured the transcript levels of 84 proinflammatory cytokines and chemokines in the livers of WT, Sod1KO and Sod1KO mice treated with Nec-1s. The data are shown as heat map in Supplementary Figure 3A, and levels of the transcripts of the proinflammatory cytokines and chemokines are presented in Supplementary Table 1. The expression of 16 of the 26 chemokines measured were significantly upregulated (Ccl1, Ccl12, Ccl17, Ccl2, Ccl20, Ccl22, Ccl3, Ccl4, Ccl5, Ccl7, Cx3cl1, Cxcl10, Cxcl16, Cxcl3, Pf4, Xcl1) in the livers of Sod1KO mice compared to WT mice (Figure 3A), and Nec-1s treatment resulted in a significant reduction in the expression of 6 of the upregulated cytokines (Ccl1, Ccl12, Ccl17, Ccl2, Ccl20, Ccl3, and Cxcl3). Of the 25 interleukins analyzed, 9 of them (IL10, IL11, IL12α IL12β, IL1β, IL1rn, IL6, and IL7) showed significant increase in the livers of Sod1KO mice compared to WT mice (Figure 3B), and Nec-1s treatment significantly reduced the expression of all of these interleukins, except IL11 and IL12α. Of the 10 TNF superfamily members, 5 showed a significant increase in the livers of Sod1KO mice compared to WT mice: Cd40lg, Ltb, TNFα, Tnfrsf11β, and Tnfsf13β(Figure 3C). Nec-1s treatment significantly reduced the expression of TNFα and Tnfsf13b. Thus, absence of *Sod1* is associated with increased expression of proinflammatory cytokines and chemokines in the liver and blocking necroptosis reduced the expression of the majority of the cytokines and chemokines, suggesting that increased necroptosis is a major contributor to increased expression of proinflammatory cytokines in livers of Sod1KO mice.

**Figure 3.**
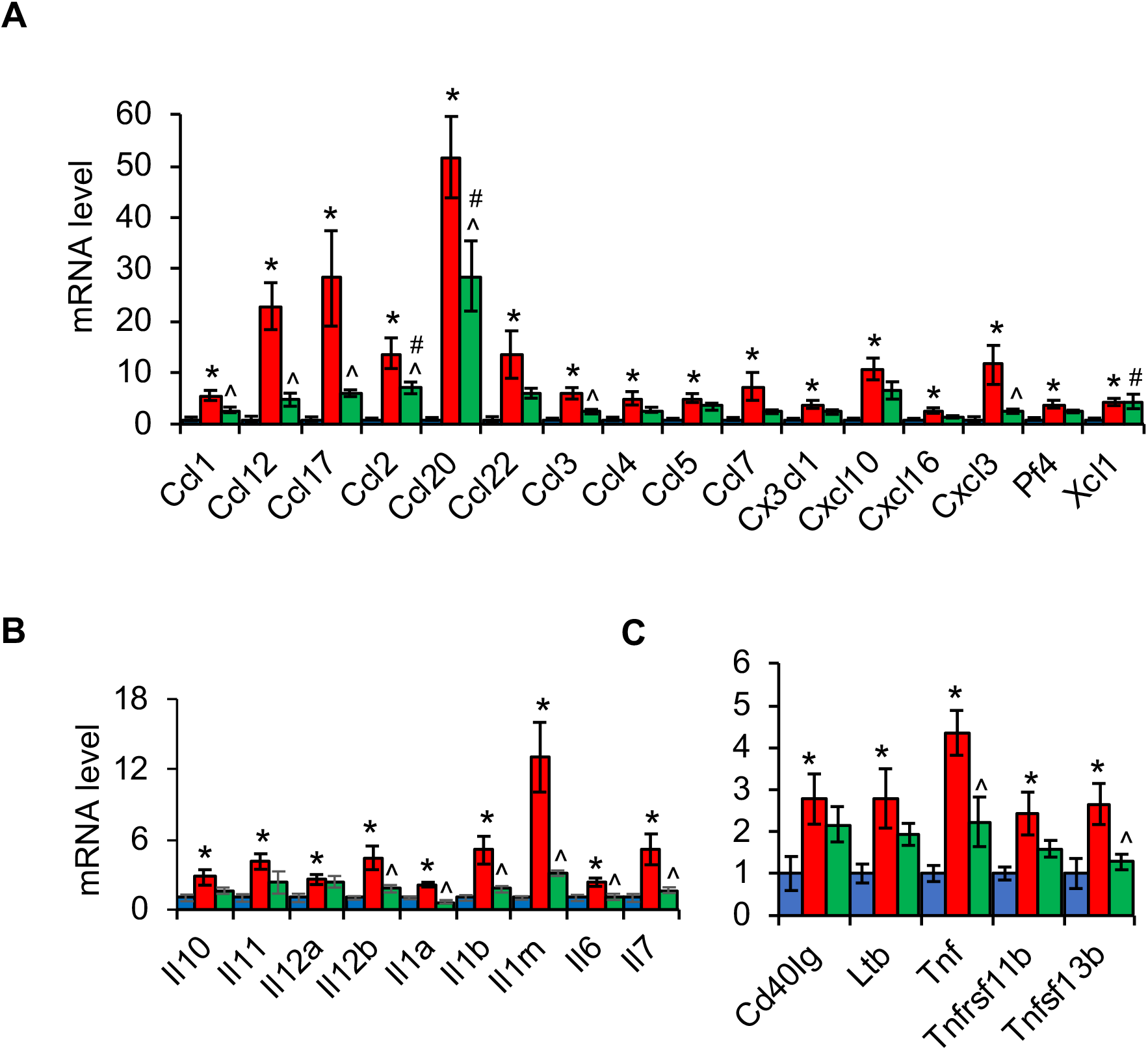
Effect of *Sod1* deficiency and necroptosis on proinflammatory cytokines and chemokines. Transcript levels of proinflammatory cytokines and chemokines **(A)**, interleukins **(B)**, and members of TNFαfamily **(C)** in the livers of WT (blue bars), Sod1KO untreated (red bars) and Sod1KO mice treated with Nec-1s (green bars). Data were obtained from 5 to 7 mice per group and are expressed as the mean + SEM. (ANOVA, *WT-Veh vs Sod1KO-Veh; ^#^WT-Veh vs Sod1KO-Veh; ^^^ Sod1KO-Veh vs Sod1KO-Nec-1s; ^*/#/^^ P ≤ 0.05).

### Effect of Sod1 deficiency and necroptosis on markers of fibrosis

The Sod1KO mice are characterized by the development of liver fibrosis, a condition that proceeds NASH (23). Because inflammation is a key driver of fibrosis in the liver (35, 36), we wanted to determine if short-term Nec-1s treatment had any effect on markers of fibrosis in the Sod1KO mice. We first measured the expression of genes associated with fibrosis in WT and Sod1KO mice treated without or with Nec-1s. The pro-fibrogenic cytokine TGFβ is produced by activated macrophages and plays a critical role in fibrosis by transforming quiescent hepatic stellate cells (HSCs) to transdifferentiated myoblasts, which produce extracellular matrix components (ECM) (37). In addition to TGFβ, pro-fibrotic cytokines connective tissue growth factor (CTGF) and platelet-derived growth factor (PDGF) are also involved in fibrosis (38). Therefore, we first tested whether expression of these pro-fibrogenic cytokines are increased in Sod1KO mice. As shown in Figure 4A, levels of TGFβ (2.2-fold), CTGF (2.6-fold), PDGFα(3.2-fold) and PDGFβ(2.7-fold) transcripts were significantly up-regulated in the livers of Sod1KO mice compared to WT mice, and Nec-1s treatment significantly reduced the expression of these pro-fibrogenic cytokines. Consistent with an increase in the transcripts of TGFβ, levels of TGFβ protein were also up-regulated in the livers of Sod1KO mice, and Nec-1s treatment reduced TGFβ protein expression in Sod1KO mice (Figure 4B). Next, we measured the expression of desmin, a protein that is strongly upregulated in liver fibrosis and is produced by activated HSCs (39). Levels of desmin protein expression were significantly upregulated in the livers of Sod1KO mice (2.1-fold) compared to WT mice (Figure 4C). Nec-1s treatment reduced levels of desmin protein by 38%, however, the decrease did not reach statistical significance. Consistent with the activation of HSCs, expression of ECM components Col1α2 (7.8-fold) and Col3α1 (7.4-fold) were significantly elevated in the livers of Sod1KO mice, and Nec-1s significantly reduced their expression (Figure 4D). In addition to ECM components, activated HSCs also express matrix metalloproteinases (MMPs) that degrade ECM, and inhibitors of metalloproteinases (TIMPs) that stabilize the ECM against MMP degradation (40). Levels of MMPs and TIMPs transcripts were significantly upregulated in Sod1KO mice liver; however, Nec-1s had no effect on the expression of these MMPs and TIMPs (Figure 4E and Supplementary Table 2). Thus, fibrosis markers are upregulated in the livers of Sod1KO mice, and short-term Nec-1s treatment attenuated expression of most of the fibrosis markers associated with HSC activation.

**Figure 4.**
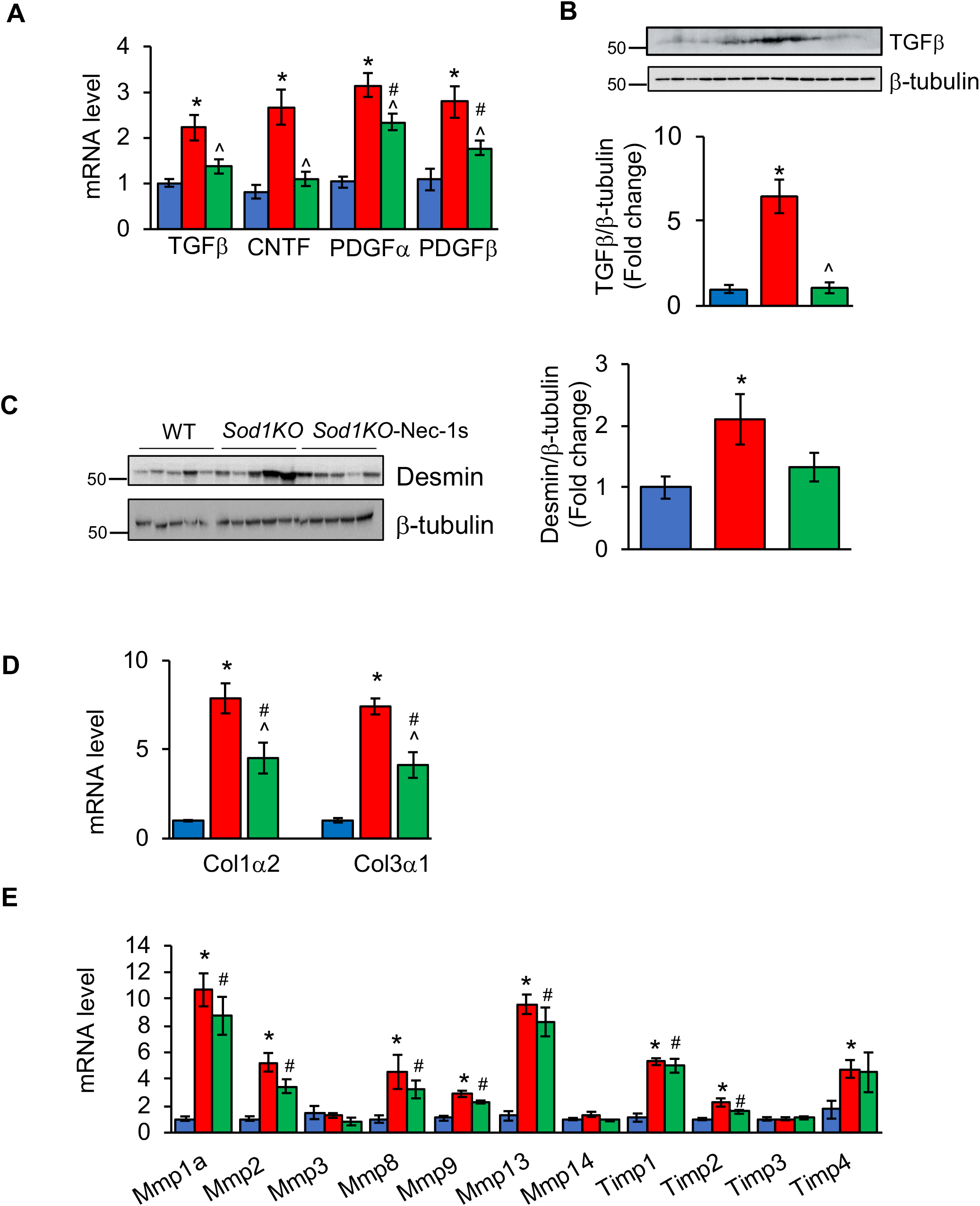
Effect of *Sod1* deficiency and necroptosis on markers of fibrosis. **(A)** Transcript levels of pro-fibrotic cytokines TGFβ, CNTF, PDGFα, and PDGFβin the livers of WT (blue bars), Sod1KO untreated (red bars) and Sod1KO mice treated with Nec-1s (green bars). **(B)** *Top panel:* Immunoblots of liver extracts for TGFβand β-tubulin. *Bottom panel*: Graphical representation of quantified blots normalized to β-tubulin. **(C)** *Left panel:* Immunoblots of liver extracts for desmin and β-tubulin. *Right panel*: Graphical representation of quantified blots normalized to β-tubulin. **(D)** Transcript levels of Col1α2 and Col3α1. **(E)** Transcript levels of MMPs and TIMPs. Data were obtained from 5 to 7 mice per group and are expressed as the mean + SEM. (ANOVA, *WT-Veh vs Sod1KO-Veh; ^#^WT-Veh vs Sod1KO-Veh; ^^^ Sod1KO-Veh vs Sod1KO-Nec-1s; ^*/#/^^ P ≤ 0.05).

### Effect of Sod1 deficiency and necroptosis on signaling pathways associated with HCC

Next, we tested whether signaling pathways that are known to modulate HCC progression and to be affected by inflammation are altered in the livers of Sod1KO mice: STAT3, JNK, ERK and p38 pathways (41–44). Activation of the oncogenic transcription factor STAT3 (P-STAT3/STAT3) was significantly upregulated in the livers of Sod1KO mice (2-fold) compared to WT mice, and Nec-1s treatment significantly reduced STAT3 activation (Figure 5A). JNK activity (P-JNK/JNK) was significantly increased in the livers of Sod1KO mice(1.7-fold). Nec-1s reduced its activity; however, this reduction did not reach statistical significance (Figure 5B). ERK activity (phospho-ERK/ERK) was similar in the livers of Sod1KO and WT mice, and Nec-1s treatment had no effect on ERK activity in Sod1KO mice (Figure 5C). The activity of p38 (P-p38/p38) was significantly lower in Sod1KO mice liver compared to WT mice, and Nec-1s treatment had no effect on p38 activity (Figure 5D).

**Figure 5.**
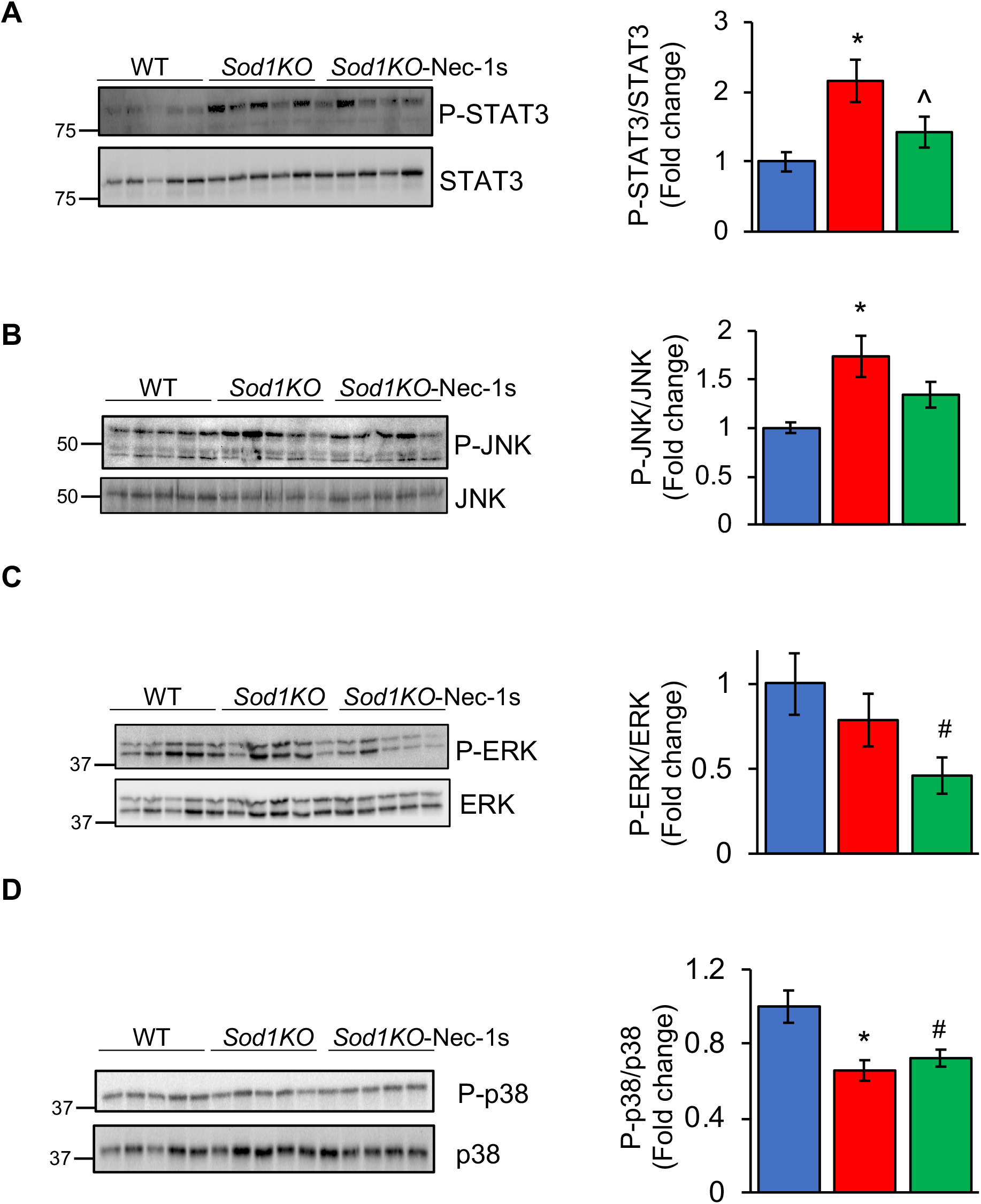
Effect of *Sod1* deficiency and necroptosis on signaling pathways associated with HCC. *Left panel:* Immunoblots of liver extracts prepared from WT (blue bars), Sod1KO untreated (red bars) and Sod1KO mice treated with Nec-1s (green bars) for P-STAT3 and STAT3 (A), P-JNK and JNK (B), P-ERK and ERK, (D) and P-p38 and p38. *Right panel*: Graphical representation of quantified phospho-blots normalized to total protein. Data were obtained from 5 to 7 mice per group and are expressed as the mean + SEM. (ANOVA, *WT-Veh vs Sod1KO-Veh; ^#^WT-Veh vs Sod1KO-Veh; ^^^ Sod1KO-Veh vs Sod1KO-Nec-1s; ^*/#/^^ P ≤ 0.05).

## Discussion

Several studies have identified non-resolving chronic inflammation as a key driver of liver fibrosis and HCC progression (48,49). Higher baseline serum levels of inflammatory markers (CRP, IL-6, C-peptide) are associated with an increased risk of developing HCC in the general population (47) and use of anti-inflammatory drugs are linked to lower risk and better survival in patients with HCC, supporting a role of chronic inflammation in HCC (48–50). Therefore, we were interested in determining if inflammation plays a role in CLD, which is observed in Sod1KO mice. Previously, we showed that circulating proinflammatory cytokines levels were increased in Sod1KO mice (51). In this study, we demonstrated that the expression of proinflammatory cytokines and chemokines, and markers of proinflammatory macrophages are dramatically increased in the livers of Sod1KO mice. For example, the expression of TNFα, IL-6, IL-1β, and Ccl2, which have been shown to increase in humans with NASH and HCC (55,56), were increased in the livers of Sod1KO mice (TNFα, 4.3-fold; IL6, 2.3-fold; IL-1β, 5-fold; and Ccl2, 13.7-fold). In addition, expression of proinflammatory M1 macrophages and levels of NLRP3 were also increased. These data are consistent with the human data showing that these measures of inflammation in the liver are associated with CLD and HCC.

Studies have shown that necroptosis is a major source of inflammation in a variety of tissues, and genetic or pharmacological inhibition of necroptosis can block/reduce inflammation (28). In addition, necroptosis has been reported to be elevated in the livers of patients with CLD (16,17) and also in the livers of mouse models of diet-induced NAFLD (15–18). Therefore, we measured necroptosis in the livers of the Sod1KO mice. We found that necroptosis was dramatically elevated in the livers of the Sod1KO mice. For example, the level of P-MLKL, the biomarker of necroptosis was increased 1.7-fold, as well as levels of MLKL (5.8-fold), Ripk3 (2.1-fold) and P-Ripk3 (2.4-fold).

Several factors have been proposed to induce necroptosis, and oxidative stress is one of them (28), e.g., deficiency of the antioxidant enzyme glutathione peroxidase 4 in hematopoietic cells resulted in increased ROS generation and necroptosis in erythroid precursor cells (54); excessive acetaminophen treatment resulted in increased ROS production and necroptosis in mice liver (55), and exposure to hypoxia increased oxidative stress and necroptotic cell death in the lung tissue of rats (56). Because of the lack of the antioxidant Cu/Zn-SOD, the Sod1KO mice show a dramatic increase in oxidative stress in various tissues including liver (26,32). Thus, our data are consistent with oxidative stress playing an important role in inducing necroptosis. In addition, our data indicate that the increased necroptosis observed in the Sod1KO mice arising from oxidative stress is correlated with increased transcription of the Mlkl and Ripk3 genes.

Because our data showed that the increase in inflammation in the livers of Sod1 mice was associated with increased necroptosis, we directly tested whether necroptosis was responsible for the increased inflammation using a RIPK1 inhibitor, Nec-1s to block necroptosis (29). Nec-1s reduced markers of necroptosis to the levels seen in wild type mice and dramatically reduced inflammation in the livers of Sod1KO mice. In particular, levels of TNFα, IL6, IL-1β and Ccl2 expression that are associated with human NASH and HCC, are significantly downregulated by Nec-1s treatment in the livers of Sod1KO mice. This reduction in inflammation in response to Nec-1s appears to arise from a decrease in M1 macrophages and reduced NLRP3 expression. Even though apoptosis was increased in Sod1KO mice liver, Nec-1s treatment had no effect on apoptosis. Thus, our data conclusively demonstrates that necroptosis contributes to increased inflammation observed in livers of Sod1KO mice.

Since chronic inflammation is believed to be a major driver of fibrosis and HCC (7, 58, 59), we were interested in determining whether blocking necroptosis and inflammation for a month had any impact on markers of fibrosis or HCC in the livers of the 10- to 13-month-old Sod1KO mice. Fibrosis markers were strongly up-regulated in the livers of Sod1KO mice, and short-term Nec-1s treatment reduced levels of fibrosis markers involved in ECM synthesis. Similarly, the increase in activated STAT3, which mediates tumor promotion (62,63), is significantly reduced by Nec-1s treatment. Thus, our preliminary data show that short-term inhibition of necroptosis, which reduces inflammation, has the potential to reduce fibrosis and HCC progression in Sod1KO mice.

In summary, our data demonstrate that increased oxidative stress in the Sod1KO mice leads to increased necroptosis and necroptosis-mediated inflammation, which appears to contribute to CLD and HCC seen in the Sod1KO mice. Because inflammation is a major factor in CLD and HCC in humans and because markers of necroptosis are increased in CLD and HCC, our data lead us to propose that necroptosis plays an important role in CLD and HCC as shown in Figure 6. We propose that obesity leading to NAFLD induces oxidative and endoplasmic reticulum stress, which triggers necroptosis in the liver leading to the generation of DAMPs. The DAMPs, in turn activate macrophages and the inflammasome leading to the production of pro-inflammatory cytokines resulting in non-resolving chronic inflammation. The increased inflammation activates HSC leading to fibrosis through increased production of ECM and eventually to HCC through the activation of various pathways, including the STAT3 pathway. Based on our model in Figure 6, we predict that blocking/reducing necroptosis would be an effective strategy for treating/preventing the development of NASH and fibrosis as well as HCC because we demonstrated that short-term (~1 month) administration of the necroptosis inhibitor, Nec-1s completely blocked necroptosis and dramatically reduced inflammation in the livers of Sod1KO mice. Currently, we are testing the ability of genetic and pharmaceutical strategies that block necroptosis to treat HCC in various mouse models, including Sod1KO mice. Several drugs are currently available that target necroptosis in addition to Nec-1s, e.g., RIPK1 inhibitor RIPA-56 (62), RIPK3 inhibitors dabrafenib (63) and GSK872 (64), and MLKL inhibitor necrosulfonamide (65). Because some of these drugs are approved for use in humans, targeting necroptosis as a potential translatable treatment for HCC in humans is potentially possible.

**Figure 6.**
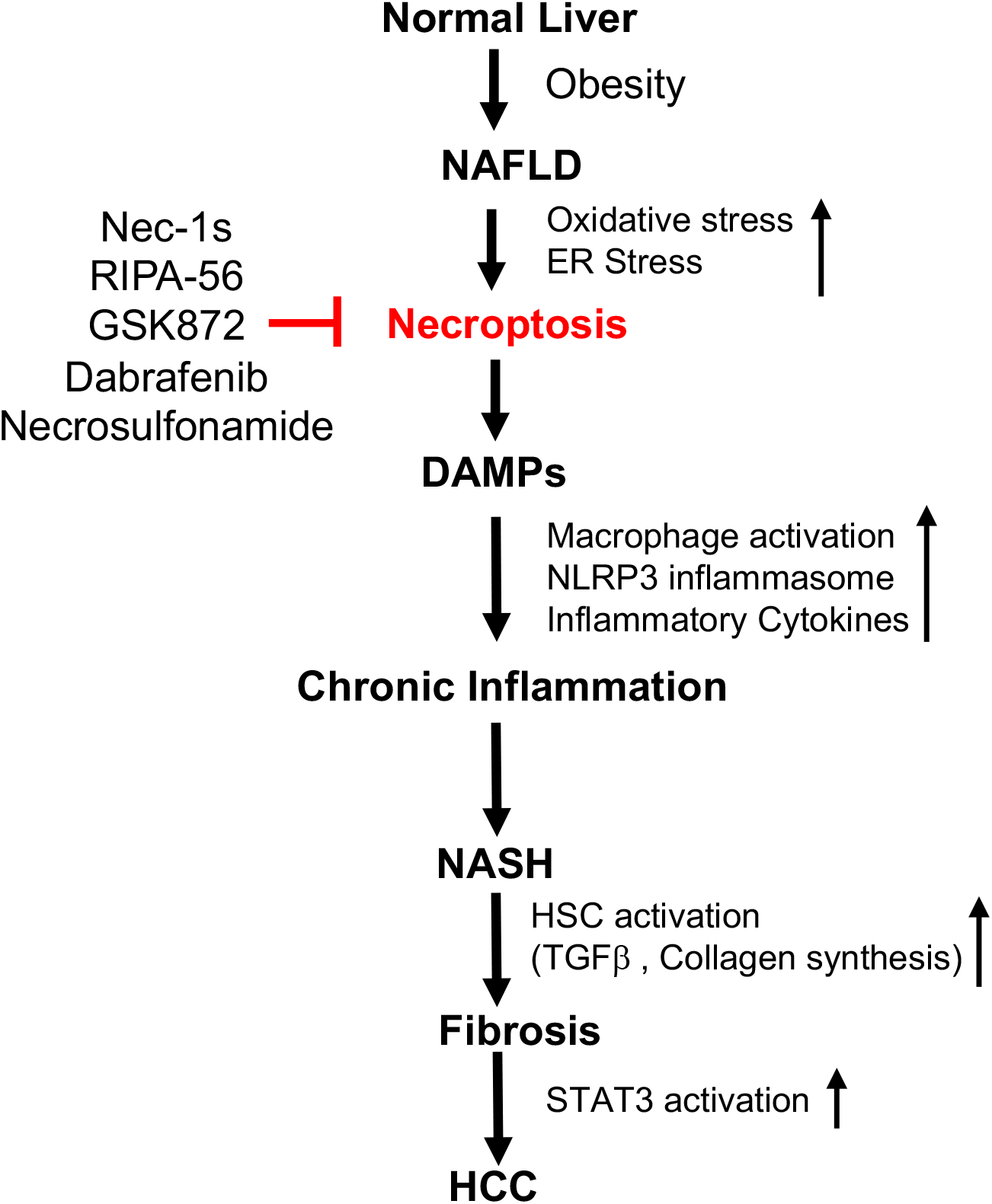
Proposed model for the role of necroptosis in CLD and HCC. NAFLD in obesity induces oxidative and endoplasmic reticulum stress, which triggers necroptosis in the liver leading to the generation of DAMPs. The DAMPs, in turn activate macrophages and the NLRP3 inflammasome leading to the production of pro-inflammatory cytokines resulting in non-resolving chronic inflammation. The increased inflammation activates hepatic stellate cells leading to fibrosis through increased production of ECM and eventually to HCC through activation of various pathways, including the STAT3 pathway. Drugs such as Nec-1s, RIPA-56 (RIPK1 inhibitor), dabrafenib and GSK872 (RIPK3 inhibitors), and necrosulfonamide (MLKL inhibitor) are shown to effectively reduce/block necroptosis and associated inflammation.

## Experimental Procedures

### Animals

All procedures were approved by the Institutional Animal Care and Use Committee at the University of Oklahoma Health Sciences Center (OUHSC). The Sod1KO mice were generated as described previously (24,32). Experimental cohorts were raised in the OUHSC Rodent facility. Ten to 13-month-old male Sod1KO mice (C57Bl/6 background) and WT controls were used for the study. The mice were group housed in ventilated cages 20 ± 2 °C, 12-h/12-h dark/light cycle and were fed rodent chow (5053 Pico Lab, Purina Mills, Richmond, IN) *ad libitum*.

### Administration of Nec-1s

Three groups of mice (10 to 13-month-old) were used for the study: wild type mice (littermates) treated with vehicle (n=6), Sod1KO mice treated with vehicle (n=7) and Sod1KO mice treated with Nec-1s (n=5). On the first day of treatment, mice were given a single intraperitoneal injection of 10mg/kg Nec-1s (7-Cl-O-Nec-1; Focus Biomolecules, Plymouth Meeting, PA) or vehicle, followed by administration of Nec-1s in drinking water for 25 days. For supplementation in drinking water, Nec-1s was first dissolved in dimethylsufoxide (DMSO, 50% w/v) and was transferred into 35% polyethylene glycol (PEG) solution and was suspended in water containing 2% sucrose (final concentration: 0.5mg/mL of Nec-1s). Daily consumption of Nec-1s based on this protocol is reported to be 2.5–5 mg/day (31).

### Quantitative real-time PCR

Total RNA was extracted using the RNeasy kit (Qiagen, Valencia, CA, USA) from 20-50 mg frozen tissues. First-strand cDNA was synthesized using a high capacity cDNA reverse transcription kit [ThermoFisher Scientific (Applied Biosystems), Waltham, MA] and quantitative real-time PCR was performed with ABI Prism using Power SYBR Green PCR Master Mix [ThermoFisher Scientific (Applied Biosystems), Waltham, MA]. PCR arrays were performed using RT^2^ Profiler PCR Arrays from Qiagen: RT^2^ Profiler™ PCR Array Mouse Cytokines & Chemokines (PAMM-150Z), RT^2^ Profiler™ PCR Array Mouse Fibrosis (PAMM-120Z). Calculations were performed by a comparative method (2^−ΔΔ*Ct*^) using β-microglobulin, actin, and 18S as controls as described previously (66).

### Western Blotting

Liver tissues collected during sacrifice were immediately frozen in liquid nitrogen and stored at −80°C until use. For western blotting, 50 mg of tissues were homogenized in extraction buffer [50 mM4-(2-hydroxyethyl)-1-piperazineethanesulfonic acid (HEPES), pH 7.6; 150 mM sodium chloride; 20 mM sodium pyrophosphate; 20 mM β-glycerophosphate; 2 mM ethylenediaminetetraacetic acid (EDTA); 1.0% Nonidet P-40; 10% glycerol; 2 mM phenylmethylsulfonyl fluoride; and protease inhibitor cocktail (GoldBio, St Louis, MO)] or RIPA lysis buffer (ThermoFisher Scientific, Waltham, MA), and western blotting was performed using 40 μg protein as previously described(67). Images were taken using a Chemidoc imager (Bio-Rad) and quantified using ImageJ software (U.S. National Institutes of Health, Bethesda, MD, USA). The following primary antibodies were used: Anti-MLKL (phospho S345) antibody and Anti-RIP3 (phospho T231+S232) antibody from Abcam (Cambridge, MA); Anti-RIPK1 (Phospho S166) antibody, Phospho-NF-κB p65 (Ser536) antibody, NF-κB p65 antibody, TGF-β antibody, STAT-3 (Phospho Tyr705) antibody, STAT-3 antibody, Phospho-SAPK/JNK (Thr183/Tyr185) antibody, SAPK/JNK antibody, Phospho p44/42 MAPK (Erk1/2) (Thr202/Tyr204) antibody, p44/42 MAPK (Erk1/2) antibody, Phospho p38 MAPK (Thr180/Tyr182) antibody, and p38 MAPK antibody from Cell Signaling Technology (Danvers, MA); Anti-RIPK1 and RIPK3 Antibodies from Novus Biologicals (Centennial, CO); Anti-MLKL antibody from Millipore Sigma (Burlington, MA); NLRP3 antibody from Adipogen (San Diego, CA); Desmin Antibody from [ThermoFisher Scientific (Invitrogen), Waltham, MA]; β-tubulin antibody and β-actin antibody from Sigma-Aldrich (St. Louis, MO). HRP-linked anti-rabbit IgG, HRP-linked anti-mouse IgG and HRP-linked anti-rat IgG from Cell Signaling Technology (Danvers, MA) were used as secondary antibodies.

### Statistical analyses

Ordinary one‐way ANOVA with Tukey’s *post hoc* test was used to analyze data.

## Data availability

The authors declare that all the data that support the findings of the study are included in the article and supporting information.

## Acknowledgments

This work was supported by NIH grants R01AG059718 (SSD) and R01AG057424 (AR), a Senior Career Research Award (AR) and a Merit grant I01BX004538 (AR) from the Department of Veterans Affairs, Oklahoma Center for the Advancement of Science and Technology research grant (HR18-053) (SSD), Presbyterian Health Foundation (OUHSC) Seed grant (SSD).

The content is solely the responsibility of the authors and does not necessarily represent the official views of the National Institutes of Health

## Abbreviations

SOD1: Cu/Zn Superoxide dismutase
HCC: Hepatocellular carcinoma
CLD: Chronic Liver Diseases
NASH: Non-alcoholic Steatohepatitis
NAFLD: Non alcoholic Fatty Liver Disease
MLKL: Mixed Lineage Kinase domain Like pseudokinase
RIPK: receptor-interacting serine/threonine-protein kinase
STAT3: Signal Transducer and Activator of Transcription 3
DAMP: Damage Associated Molecular Patterns
NLRP3: NOD-, LRR- and pyrin domain-containing protein 3
CCL: Chemokine (C-C motif) ligand
CXCL: C-X-C Motif Chemokine Ligand
IL: Interleukin
TGF: Transforming Growth Factor
TNF: Tumor Necrosis Factor
JNK: c-Jun N-terminal kinase
ERK: extracellular-signal-regulated kinase
HSC: Hepatic stellate cells

